## Supplementary Information for "Single-molecule analysis of the entire perfringolysin O pore formation pathway"

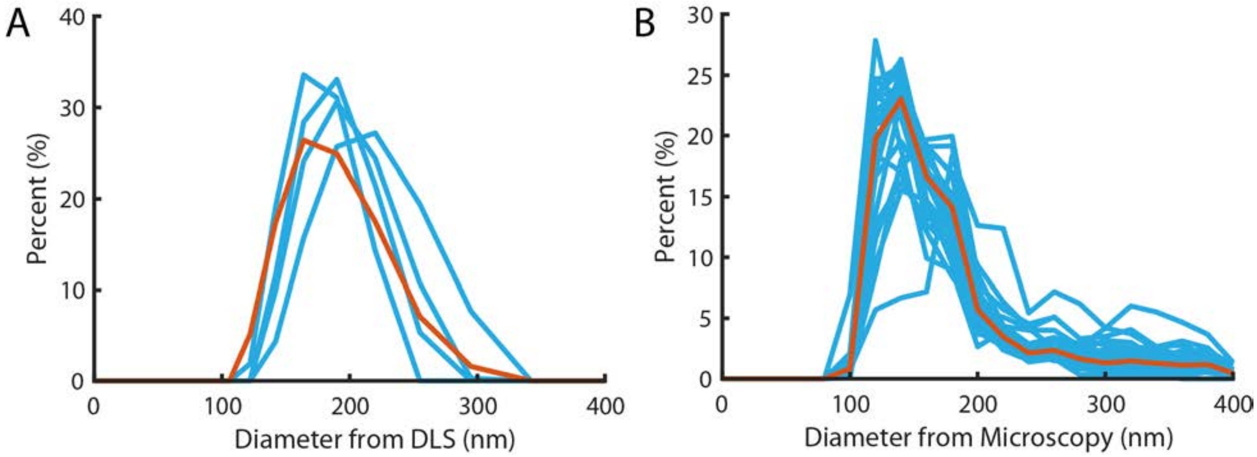


**Figure 1–Figure Supplement 1. Liposome size distribution. (A)** Size distribution measured using dynamic light scattering with a mean and standard deviation of 183±37 nm. Blue lines denote the four repeats, red line is the pooled distribution. **(B)** Size distribution measured from diameters of particles in super-resolution reconstructions (Figure 2G). Light Blue lines denote the distribution of independent experiments, the red line is the pooled distribution with a mean and standard deviation of 172±18 nm.

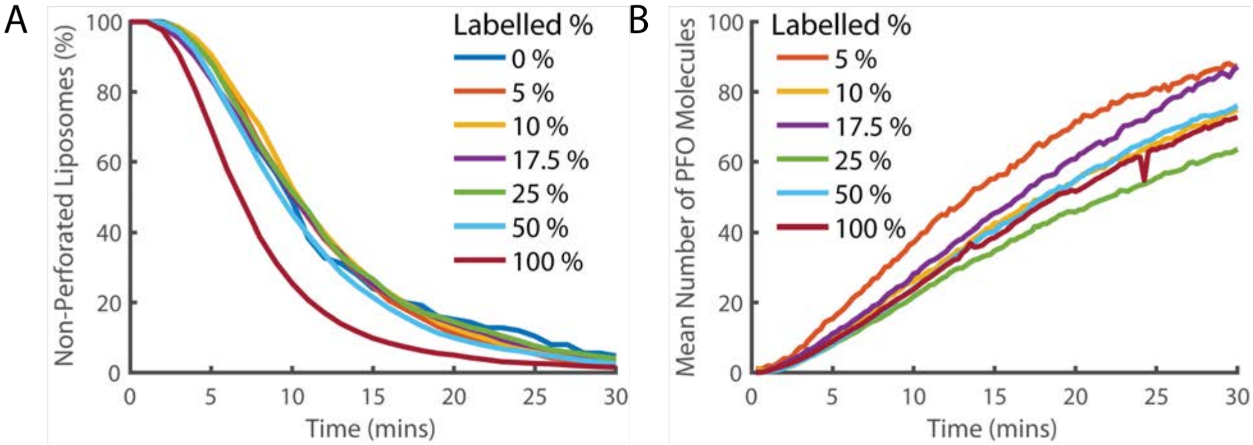


**Figure 1–Figure Supplement 2. Fluorescence labelling of PFO does not impair pore formation activity. (A)** Distributions of single-step dye release times from liposomes in the presence of different ratios of labelled and unlabelled PFO (combined concentration of 400 pM). The percentage denotes the percent of the 400 pM PFO that is fluorescently labelled by AF647. Concentrations of labelled and unlabelled protein were independently measured by gel densitometry. The slight increase in the rate of poration in the 100% labelled sample (maroon curve) in comparison to lower labelled percentage samples suggests that the actual concentration of the labelled sample may be slightly higher than the unlabelled sample. Alternatively, this could be the result of fluorescent labelling increasing the activity of PFO. Regardless, the activity of all samples is reasonably similar and we conclude that the labelled PFO is functional. **(B)** The average across all liposomes of the PFO intensity over time. The intensity is corrected for the partial labelling of PFO by dividing by the labelling percent. If PFO does not experience fluorescence quenching when it forms an oligomer, then all traces should overlay. The relatively small deviation between traces and absence of a clear trend with concentration suggests that the degree of quenching is small.

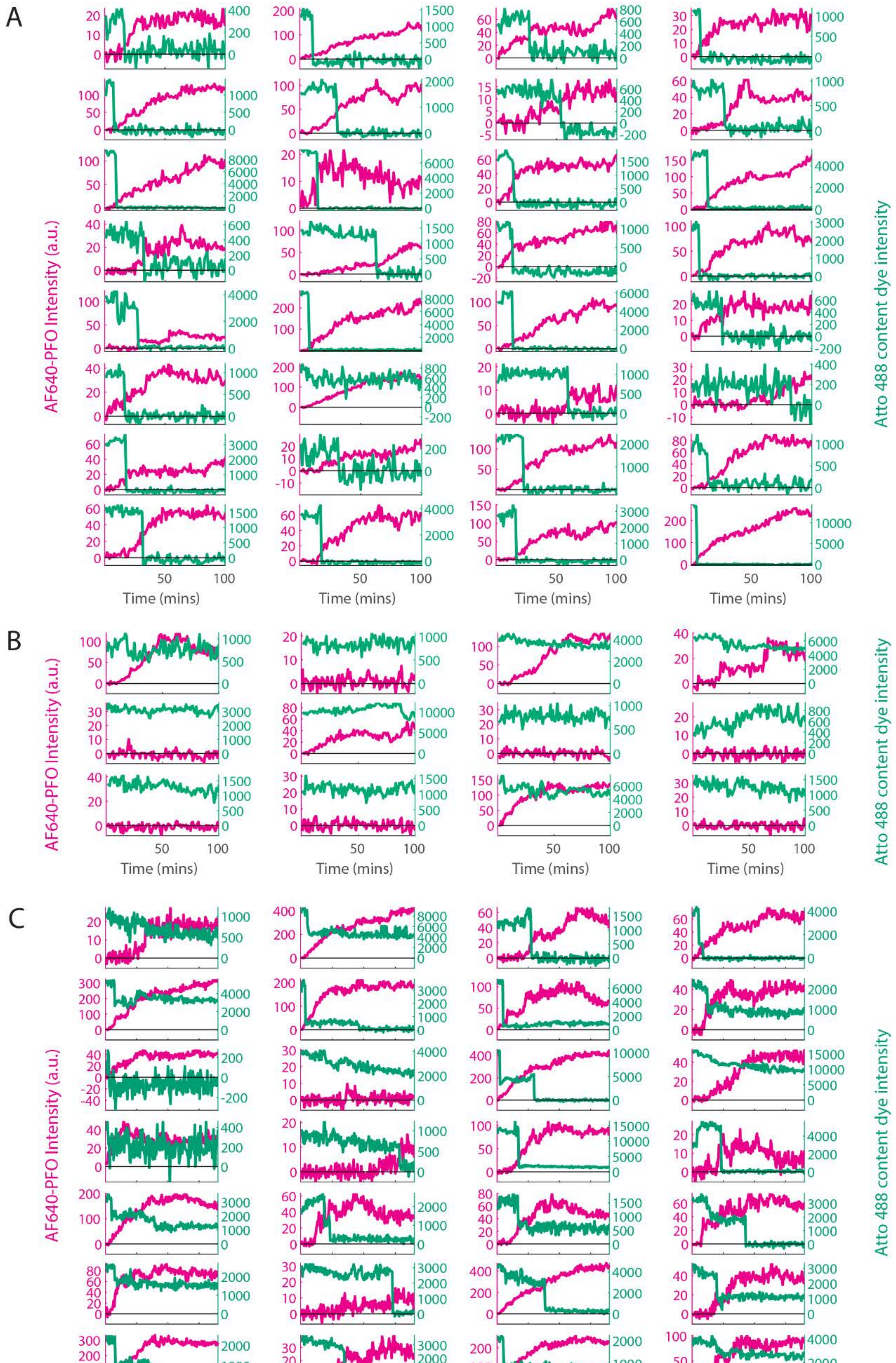
A
B
C
AF640-PFO Intensity (a.u.)
Atto 488 content dye intensity
Time (mins)

**Figure 1–Figure Supplement 3. Classification of single-liposome traces on the basis of the content dye release profile.** Content dye (blue-green) and corresponding AF647-PFO (magenta) binding traces from a dual colour TIRF pore formation experiment recorded in the presence of 100 pM AF647-PFO. The traces were classified on the basis of step-fitting of content dye traces as (A) single-step dye release, (B) no dye release (defined as less than 25% decay in content dye intensity over the course of the experiment), (C) other dye release profiles (includes multi-step, partial dye and gradual dye release). Categories B and C were excluded from analysis. After poration, liposomes continue to bind AF647-PFO to levels greater than the value expected for a single ring-shaped pore (~35 subunits), suggesting that eventually multiple pores form on a single liposome. The example traces shown here represent the first 32 out of 457, 12 out of 195 and 28 out of 280 traces in category A, B and C, respectively for this experiment (i.e. these traces were not hand-picked).

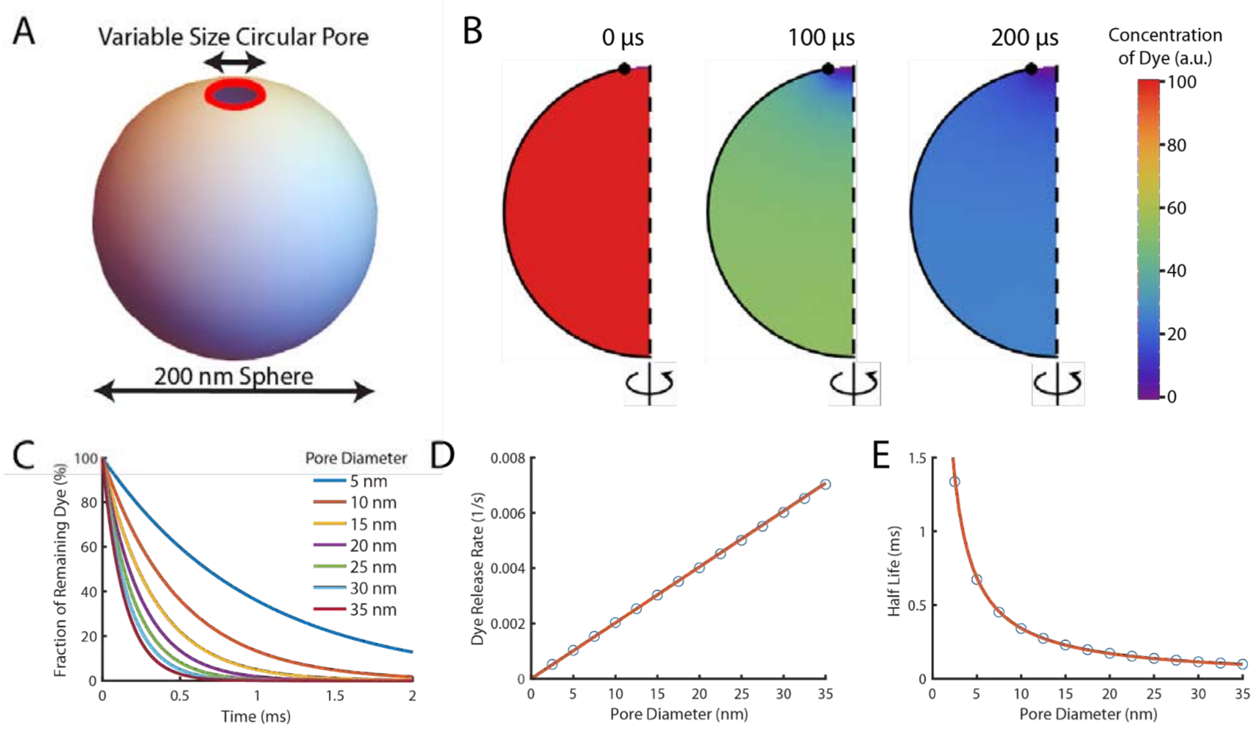


**Figure 1–Figure Supplement 4. Theoretical computation of the dye release rate following pore opening.** Pore formation on liposomes was modelled in FlexPDE v7 as a circular pore on a spherical liposome. Encapsulated dye was initially homogeneously distributed in the liposome before being allowed to diffuse out through the pore at a rate of 435 $\mu m^2/s$ (Petrásek and Schwille 2008) . **(A)** Schematic of the simulated system. The liposome was modelled as a sphere with a diameter of 200 nm, and the pore (shown in red) was assumed to be a ring with a variable diameter to model different size pores **(B)** A montage of the distribution of dye inside a liposome with a 35 nm pore. Each semicircle is a cross-section of the liposome at a given time point which would be rotated around the z-axis to give the 3-dimensional model. The pore is at the top of the liposome (the lack of black outline at the top of the semicircle). Initially (left-hand panel), the concentration is homogeneously at 100%. After 100 µs (middle panel) approximately half the dye has left the liposome. The concentration is lowest near the pore (indicated by the blue colour at the top). After 200 µs (right panel) little dye remains in the liposome. **(C)** The fraction of dye remaining in liposomes with various pore sizes as a function of time. Concentrations exponentially decay, with dye escaping faster from liposomes with larger pores (the maroon line tends to zero faster than the dark blue line). **(D)** The rate of dye release, calculated by exponentially fitting curves from (C), increases linearly with pore size. The linear fit of the rate of dye release distribution (orange line) gives a dye release rate constant of 2.02 $nm^{-1}$ $s^{-1}$. **(E)** Plot of the corresponding half-lives of dye release.

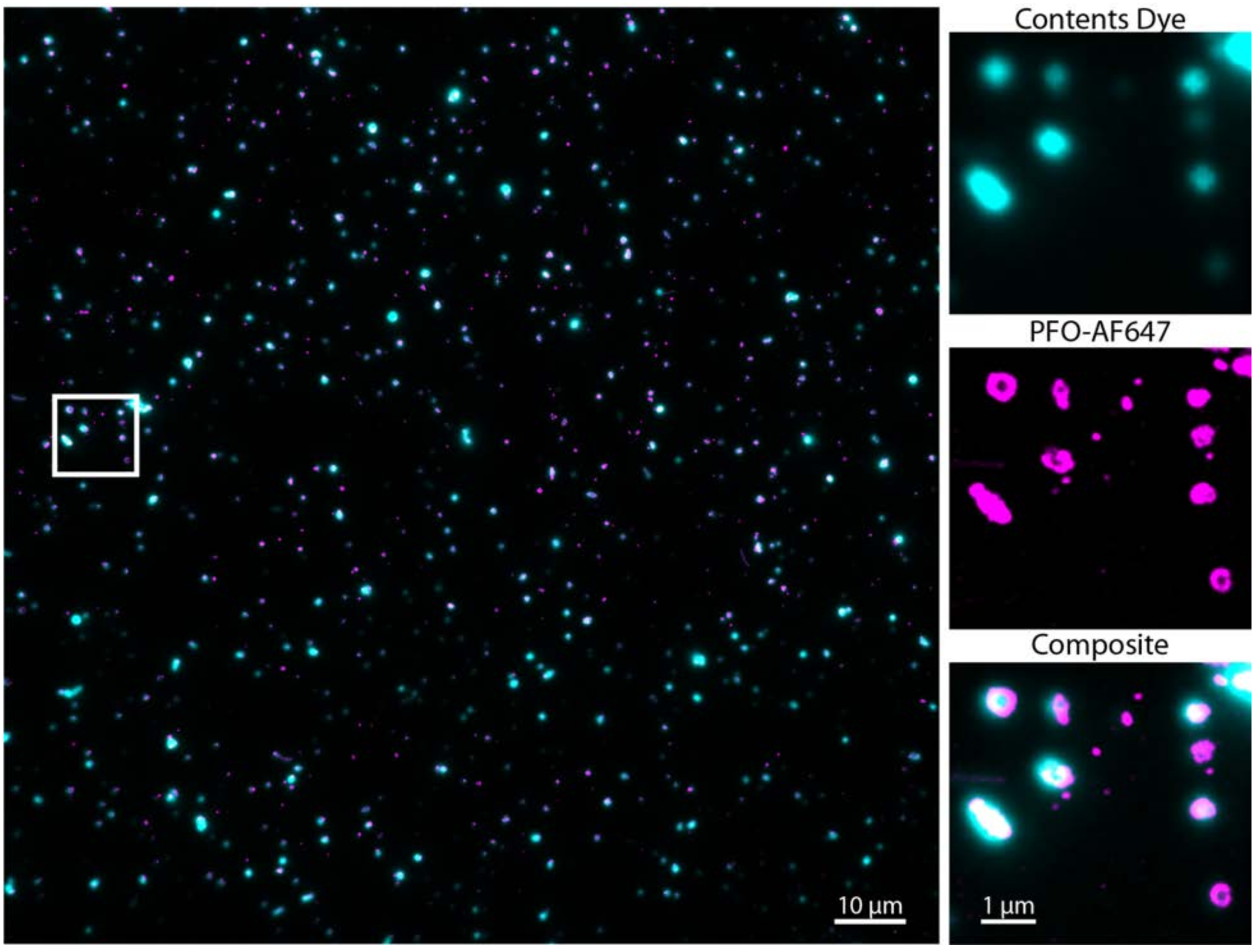


**Figure 2–Figure Supplement 1. Single-molecule AF647-PFO binding events co-localise with Alexa Fluor 488-loaded liposomes.** Overlay of a super-resolved AF647-PFO image reconstructed from single-molecule localisations (magenta) and the corresponding TIRF image of Alexa Fluor 488-loaded liposomes (cyan); full field of view (left) and zoomed-in region (right) contained within the white box (corresponding to the region in Figure 2G/H). Localisations are determined by point-spread function fitting of AF647-PFO spots in each frame of a TIRF stack with 432000 frames acquired at a frame rate of ~17.5 frames/s in the presence of 10 pM AF647-PFO.

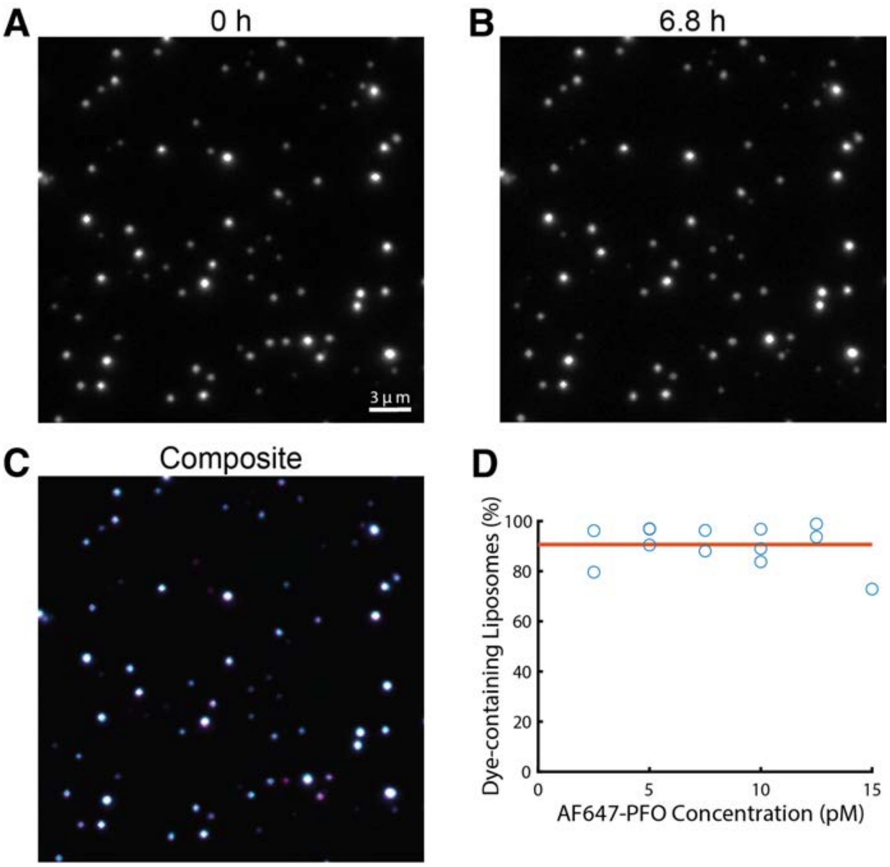


**Figure 2–Figure Supplement 2. Liposomes retain their content dye during single-molecule PFO binding experiments. (A-C)** TIRF images from a time series recorded in the presence of 20 pM AF647-PFO at the beginning (A, 0 h) and the end (B, 6.8 h) of a single-molecule PFO binding experiment. The overlay (C) of both images shows that most liposomes retained the content dye during exposure to low concentrations of PFO for 6.8 h. **(D)** Plot of the fraction of dye-containing liposomes remaining at the end of the single-molecule PFO binding experiments (2.5–15 pM AF647-PFO) with fit line (orange) showing that on average  91% of liposomes remain unperforated over the course of the experiment.

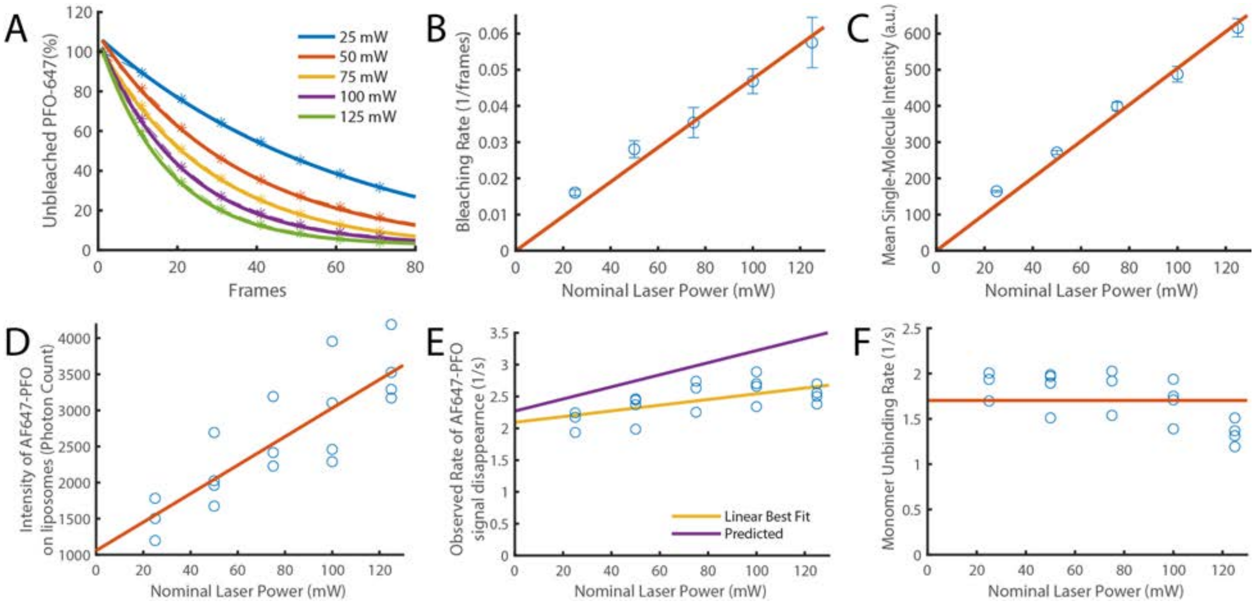


**Figure 2–Figure Supplement 3. Validation of photobleaching correction of single-molecule unbinding rates. (A-C)** Single-molecule photobleaching experiments of AF647-PFO adsorbed onto the surface of a glass coverslip were performed at nominal laser powers between 25–125 mW and analysed using JIM. (A) AF647-PFO photobleaching curves with exponential fits to determine the bleaching rate at each laser power. (B) The bleaching rate increases linearly with laser power. The linear fit gives a bleaching rate constant of 4.75±0.8 x $10^{-4}$ $mW^{-1}$ $frame^{-1}$. (C) The intensity of molecules also increases linearly with laser power. Together these results confirm that the light dose for excitation of molecules increases linearly with nominal laser power. **(D-F)** Single-molecule binding experiments in the presence of 20 pM AF647-PFO were performed across the same range of laser powers. (D) The mean intensity of AF647-PFO signals on liposomes quantified using Picasso (expressed in photon counts) increases linearly with laser power. (E) The observed rate of signal disappearance of liposome-bound AF647 monomers is the combination of monomer unbinding and photobleaching and, as expected, increases linearly with the nominal laser power. The yellow line denotes the linear best fit, whereby the y-intercept is the monomer unbinding rate (2.1±0.2 $s^{-1}$) and the gradient is the bleaching rate (4.5±3.7 x $10^{-3}$ $mW^{-1}$ $s^{-1}$ or 2.25±1.9 x $10^{-4}$ $mW^{-1}$ $frame^{-1}$). The purple line represents the predicted power dependence based on the monomer unbinding rate from Figure 2N (2.27 $s^{-1}$) and the photobleaching rate calculated in panel B. (F) Data points from panel E after applying a photobleaching correction using the rate from panel B. The monomer unbinding rate is expected to be independent of the laser power. The red line denotes the mean of 1.7±0.3 $s^{-1}$.

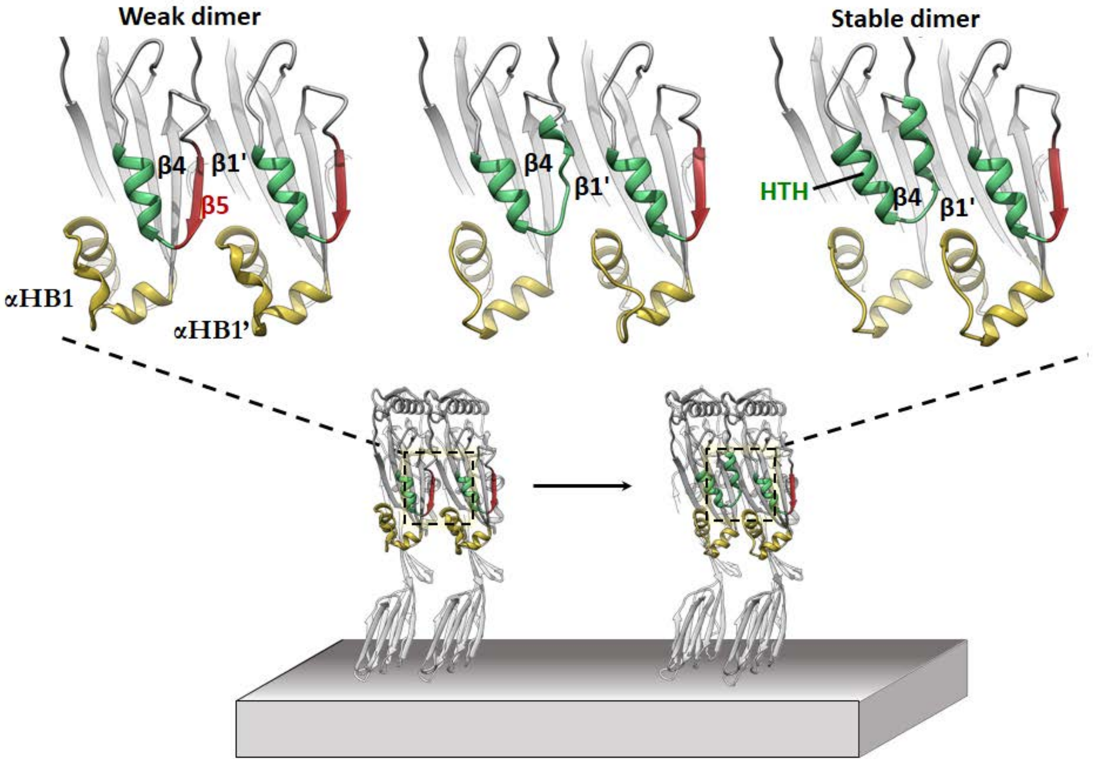


**Figure 2–Figure Supplement 4. *In silico* modelling of early-stage PFO assembly and conversion to irreversible dimer (nucleation).** AlphaFold predictions of the PFO dimer interface suggest two conformational states exist. These are consistent with a weak transient dimer which ultimately causes strand β5 to move upon binding. The dimer transitions into a stable irreversible dimer where β5 has interconverted into the HTH motif and β4-β1' hydrogen bonding defines the upper region of the nascent MACPF/CDC giant β-barrel. The predicted transient dimer formed before movement of strand β5 is reminiscent of the CDC dimers proposed on the basis of linear CDC oligomers (Lawrence et al. 2015, 2016).

**Figure 2–Figure Supplement 5. Morphing movie of the two dimer conformational states predicted by AlphaFold.**

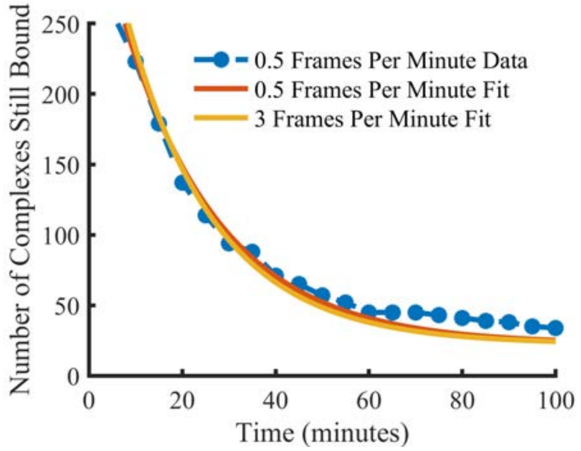


**Figure 3–Figure Supplement 1. Comparison of state lifetimes for experiments run at 3 frames per minute and 0.5 frames per minute.** Decreasing the frame rate further to 0.5 frames per minute gave a release rate of 0.048 $min^{-1}$ for the observed complex. For comparison, the 3 frames per minute release rate after photobleaching correction (0.052 $min^{-1}$) is also shown. The exponential fit had a positive offset suggesting the existence of an even longer-lived state but it is incredibly rare (~6.4% of long-lived localisations) and so was in too few numbers to be meaningfully quantified.

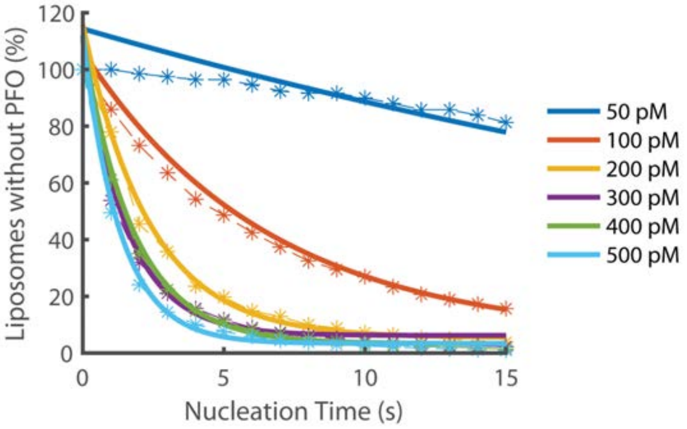


**Figure 4–Figure Supplement 1. Exponential fits of nucleation rates.** Exponential decay functions (solid lines) were fitted to the experimental data to determine the nucleation rate at each concentration.

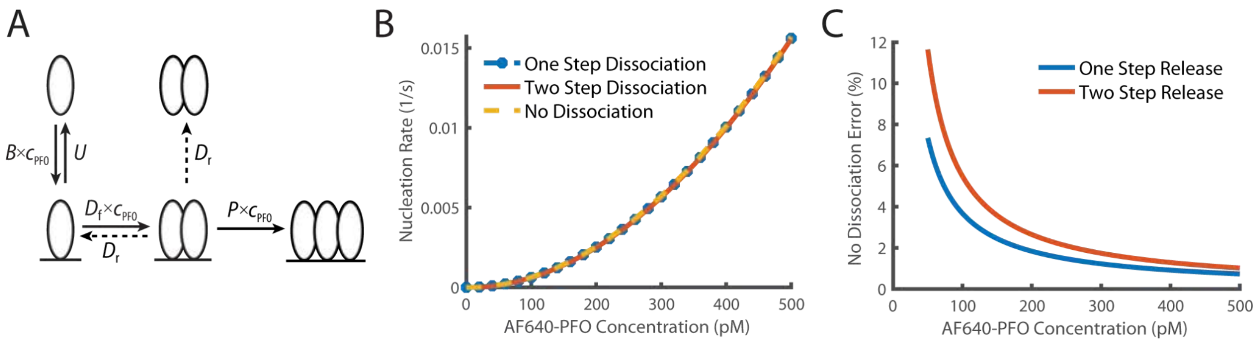


**Figure 4–Figure Supplement 2. Comparison of dimer release models. (A)** Schematic demonstrating the two potential methods of dimer dissociation, the dimer falling directly off the membrane in one step, or a two-step process, where the dimer falls apart into two membrane monomers which can then individually dissociate. **(B)** Overlay of the predicted nucleation rate based on whether it is assumed that the dimer can dissociate as a one-step, two-step process or if it is assumed that the dimer does not dissociate. **(C)** The percentage difference between no dimer dissociation and the two other models. The difference is always less than 5% for values above 100 pM.

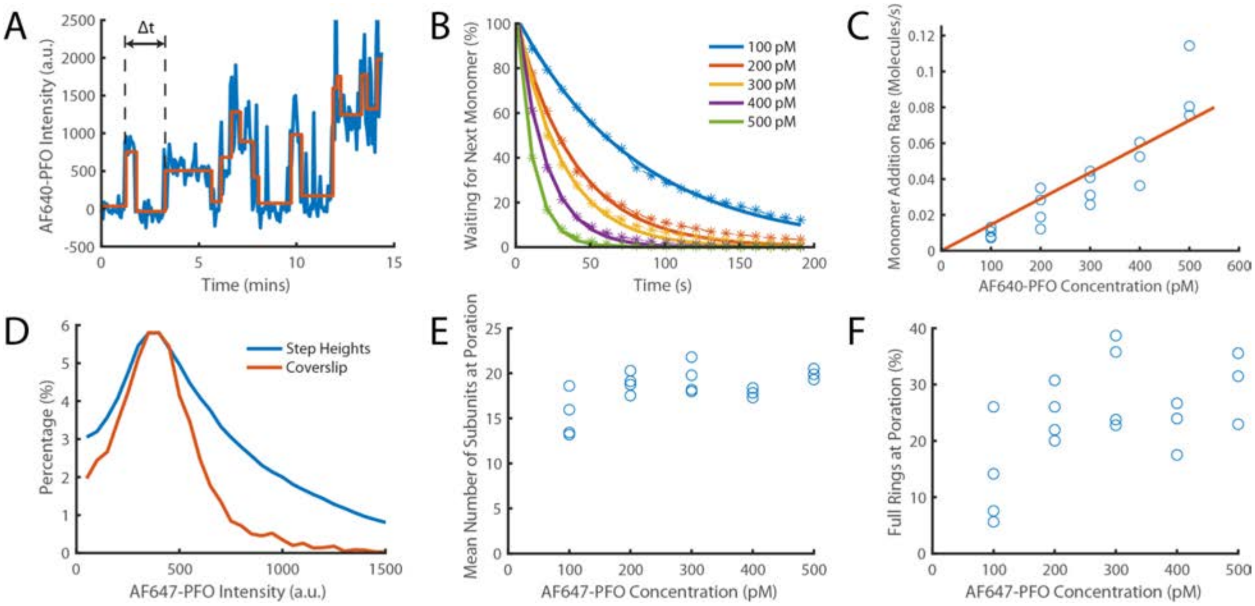


**Figure 4–Figure Supplement 3. Direct measurement of PFO monomer addition using step fitting.** The PFO assembly and pore formation (dye release) dual colour TIRF microscopy assay using dye-loaded liposomes was carried out in the presence of 100–500 pM AF647-PFO at high temporal resolution (frame rate; exposure time 50ms) and with high 640 nm laser power (75 mW) to directly image the addition of AF647-PFO monomers to a membrane-bound oligomer on liposomes. **(A)** Example AF647-PFO intensity trace (blue) up to the time of dye release (dye intensity trace not shown). The addition of an AF647-PFO monomer to the membrane-bound oligomer is observed as a stepwise increase in intensity. The higher light dose (high laser power and high temporal resolution) required for single-molecule detection leads to photobleaching, observed as negative steps in the trace. The step fit of the trace using change point analysis is shown in orange. The waiting times between monomer addition events are determined as the time differences between successive positive steps (the first waiting time in the trace is labelled Δ*t*). **(B)** Distributions of waiting times between positive steps recorded at AF647-PFO concentrations between 100–500 pM with exponential fits (solid lines). The exponent of each fit gives the rate of monomer addition for that concentration. **(C)** Plot of the monomer addition rate as a function of AF647-PFO concentration obtained from panel B. The linear fit line (orange) gave a monomer addition rate of 0.15±0.04 $nM^{-1}$ $s^{-1}$. The monomer addition rate determined here by step fitting represents an independent measurement of the oligomerisation rate (0.23±0.028, Figure 4C), whereby the lower value (by 35%) may be in part due to undetected short lived steps. **(D)** Distribution of positive step heights from all experiments compared to the intensity of single molecules bound to a coverslip. Both peaks occur at the same location, however, the step height distribution is broader (reflecting the increase in shot noise from multiple molecules) and has a longer tail in part due to missing short-lived steps. These missing steps are likely to account for the small reduction in oligomerisation rate measured in panel C compared to that measured in Figure 4C. **(E)** Mean number of positive steps (each corresponding to the addition of a monomer) before contents dye release. This analysis provides an intensity-independent estimate of the number of subunits in the oligomer at the time of membrane insertion (poration). The values plateau at similar values to Figure 6C (here ~18 versus ~21). **(F)** The percentage of full rings at poration, calculated using the number of liposomes with at least 35 positive steps before contents dye release. At high concentrations (≥200 nM) ~25% of oligomers contain the number of PFO molecules required to form a closed ring, similar to the observations in Figure 6D. Four independent repeats were performed for 100 pM, 200 pM and 300 pM and 3 repeats for 400 pM and 500 pM.

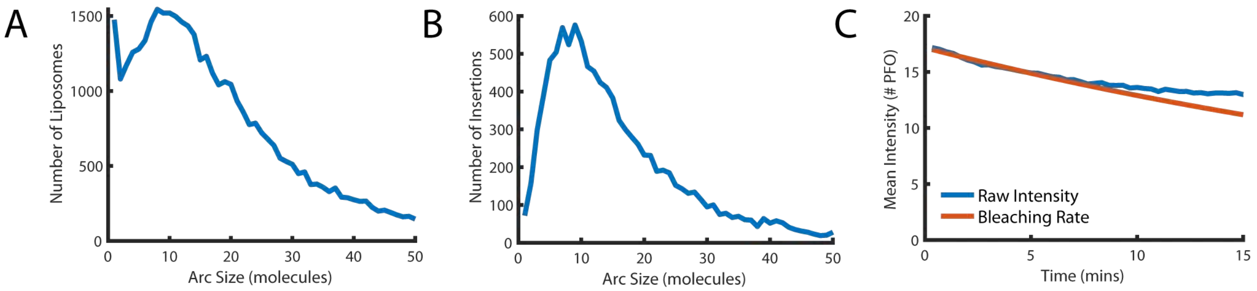


**Figure 5–Figure Supplement 1. Number of subunits in the PFO oligomer during washout. (A)** Distribution of the number of liposome-bound PFO molecules determined for all liposomes in the field of view. **(B)** Distribution of the number of subunits in PFO oligomers that proceed to insert into the membrane after AF647-PFO wash-out from solution. **(C)** Mean intensity of AF647-PFO oligomers bound to liposomes in the field of view as a function of time after wash-out of AF647-PFO from solution (blue line). The decay over time of the mean AF647-PFO oligomer intensity does not exceed the decay expected for photobleaching (red line).

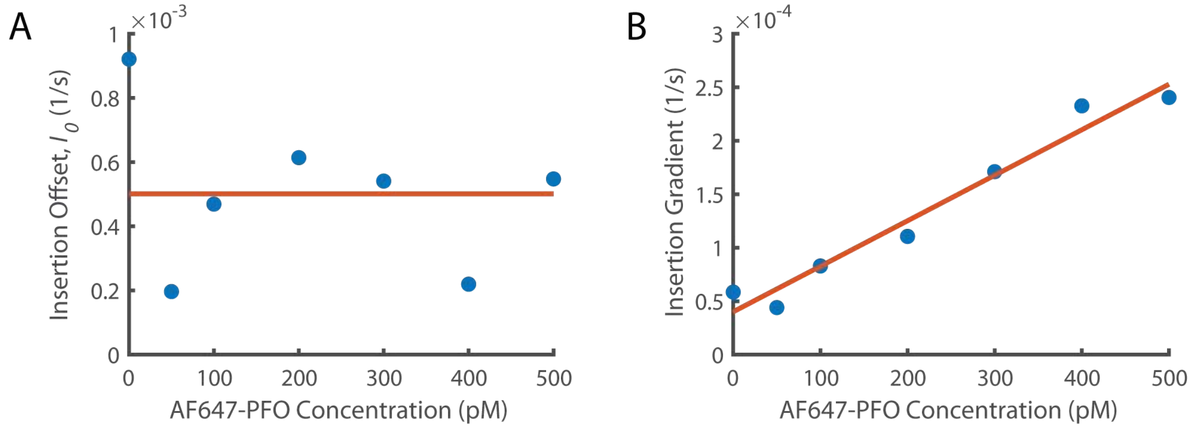


**Figure 6–Figure Supplement 1. Fitting Concentration Dependence of Insertion (A)** The insertion offset is the y intercept of all linear fits in Figure 6A. The y intercept from the washout experiments (Figure 5C) is included as the point at 0 pM. These data points are fit with a constant since there is no obvious trend. **(B)** The change in gradient for the lines of best fit from Figure 6A as a function of concentration. We observe that the gradient increases linearly suggesting that the rate of increase of insertion with the number of subunits in the prepore increases with PFO concentration. The y-intercept of the linear fit (orange line at 0 pM concentration) gives the insertion constant ($I_{g0}$) the rate of increase in insertion rate with the number of subunits in the prepore in the absence of PFO. The gradient of this fit ($I_{gc}$) is the increase of the dependence on the number of subunits in the prepore with PFO concentration.

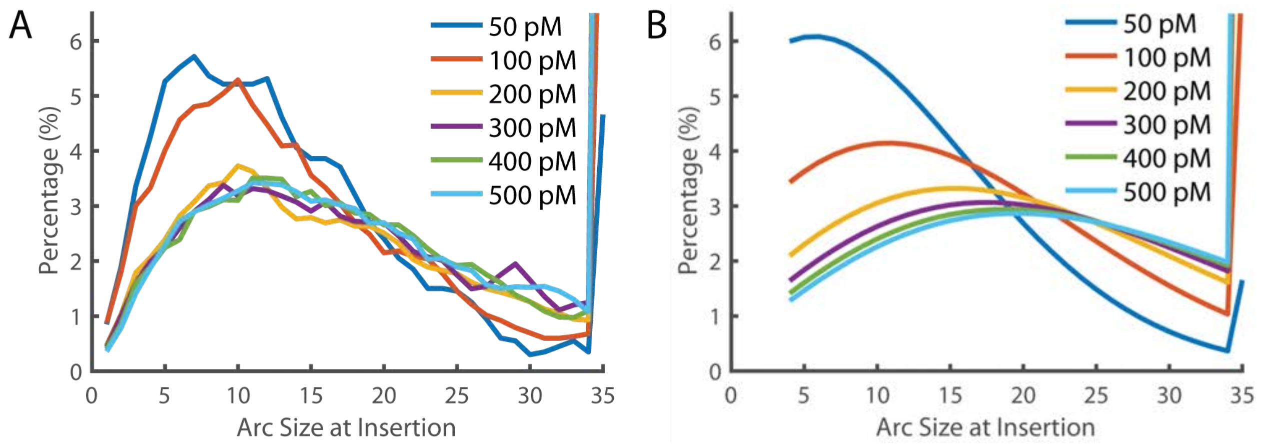


**Figure 6–Figure Supplement 2. Arc size distribution at the time of Insertion. (A)** The experimentally measured distribution of arc sizes at the time of insertion. **(B)** The theoretical prediction of arc size distribution. At low concentrations, the distribution closely reflects the experimental observation. However, at higher concentrations, a greater number of subunits in the prepore were predicted than observed. A 35-mer is considered to be a full ring that cannot add additional subunits (hence these species accumulate). See also Figure 6D for the fraction of complete rings formed at different PFO concentrations.

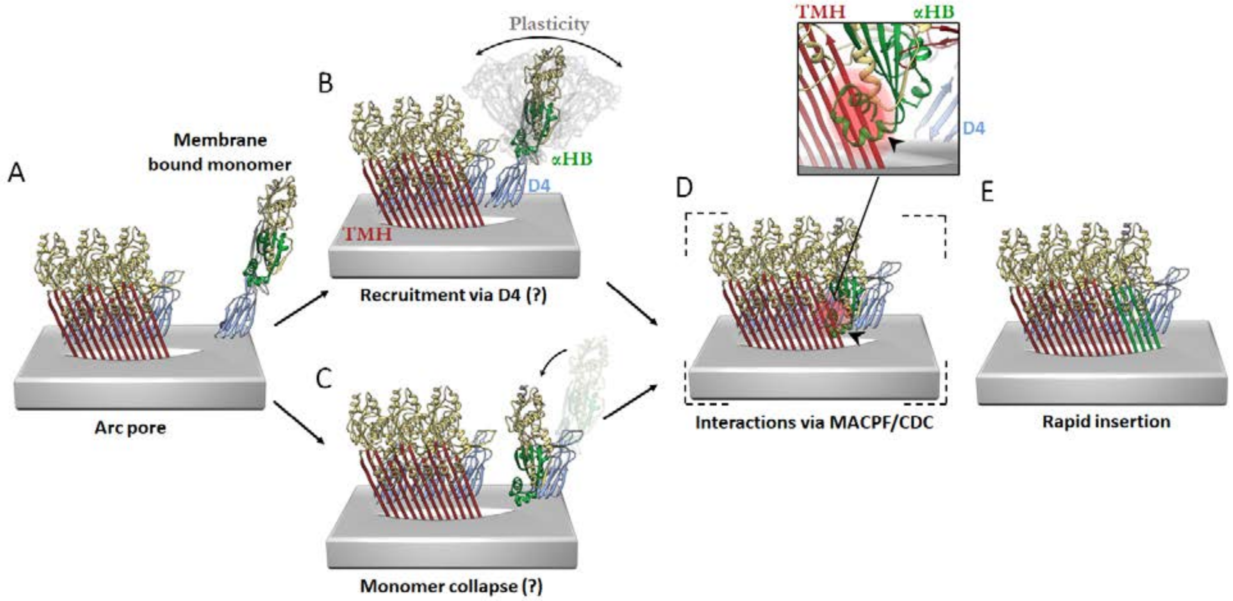


**Figure 6–Figure Supplement 3. Hypothetical models of PFO pore growth for an inserted arc pore.** **(A)** An arc of inserted PFO (trans-membrane hairpin (TMH), red; D4, blue; D1/D2/D3, yellow; three PFO molecules shown for simplicity) recruits a monomeric PFO. In order for the β-barrel to form on a growing arc, the newly recruited monomer must undergo a vertical collapse, release of the α-helical bundles and unfurling of the β-hairpins. In the canonical prepore model (not shown), these processes are thought to be driven by interactions between the MACPF/CDC domains (consisting of D1 and D3) of adjacent subunits. In contrast, the interaction interfaces involved in the recruitment and conformational changes of a membrane-bound monomer for post-insertion arc growth are unknown, but may involve transient D4–D4 interactions, which have been postulated to form early in PFO assembly (van Pee et al. 2017) and/or some of the interactions in D1 and D3 as seen in the linear oligomer structures (Lawrence et al. 2015, 2016). These possible models are described as follows. **(B)** Monomeric PFO is flexible about the hinge region between D4 and the rest of the PFO molecule (as shown in molecular dynamics simulations (Reboul, Whisstock, and Dunstone 2014)). This flexibility enables monomer collapse after recruitment. **(C)** Alternatively, membrane bound monomers are capable of collapse prior to recruitment, thereby positioning the MACPF/CDC domain appropriately for direct interactions at the MACPF/CDC interface. The trigger for PFP collapse in the alternative pathways shown in B and C are unknown, but may involve the local environment of the toroidal pore. **(D)** Subsequent more substantive interactions may then be mediated by interdomain contacts of the MACPF/CDC interface. Therefore, a putative collapsed intermediate is postulated to exist whereby collapse precedes unfurling of the α-helical bundles (α-HB) required to form transmembrane hairpins (TMH). Simple rigid body fit of the PFO crystal structure (PDB 1PFO) into the cryoEM reconstruction of the pneumolysin pore (EMD-4118) reveals a sterically forbidden clash of α-HB1 (black arrow/red glow) occurs with strand β4 of the adjacent PFO (inset). This model suggests any conformational rearrangement of α-HB1/2 must occur shortly after or during collapse. **(E)** Full release of α-HB1 and α-HB2 results in the addition of a new inserted PFO monomer.

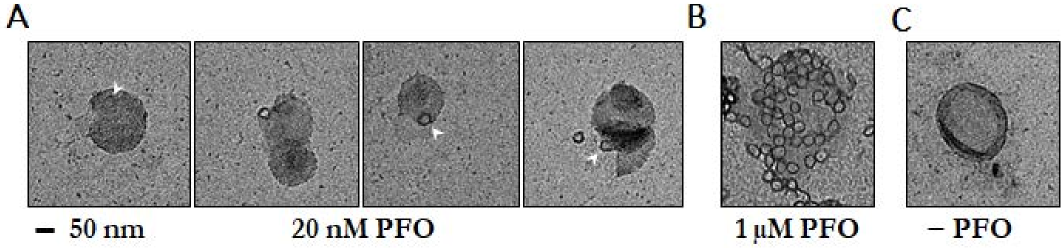


**Figure 6–Figure Supplement 4. Negative-staining electron microscopy of PFO on POPC/cholesterol liposomes. (A)** Liposomes were incubated with 20 nM PFO for 30 mins at room temperature in solution before deposition onto grids. Under these conditions, PFO oligomers were rare and appeared as single complete rings. No structures were observed on liposomes prepared with PFO concentrations in the pM range. PFO assembly on liposomes in solution for EM is likely to require higher concentrations than PFO assembly on surface-bound liposomes in the TIRF assay because of PFO depletion from solution in the former but not the latter experiment. We note that even though PFO is constantly resupplied by the microfluidic flow in our TIRF assay, measurements in the pM range require careful passivation of surfaces to prevent PFO loss due to non-specific adsorption, which can lead to failure of PFO assembly. **(B)** Liposomes incubated with 1 μM PFO show numerous rings per liposome. **(C)** Negative control, no PFO added. Scale bar; 50 nm.

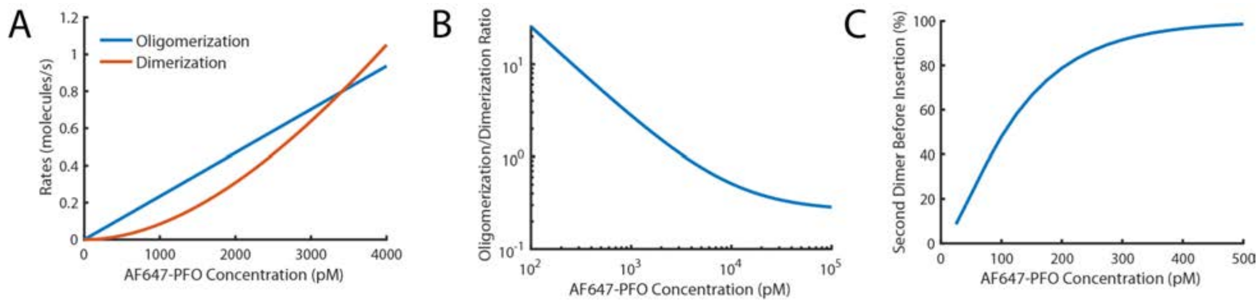


**Figure 6–Figure Supplement 5. Probability of nucleating a second PFO oligomer on the liposome before the first oligomer has inserted into the membrane. (A,B)** Fits for oligomerisation and dimerisation rates extrapolated out to a PFO concentration of 4 nM (A) and log-log plot of the ratio of the oligomerisation rate to the dimerisation rate over a range from 100 pM to 100 nM (B). The plots show that monomer addition via oligomerisation is >20-fold faster than dimer formation at 100 pM, rates for both processes are equal at 3.4 nM and oligomerisation is 5-fold slower than dimerisation at 100 nM. **(C)** Probability of forming a second PFO dimer (as a nucleus for an independently growing oligomer) on the liposome before the first oligomer has inserted into the membrane. This probability is calculated using the dimerisation rate and the nucleation to insertion time (calculated in Figure 6E). In the low concentration range used in this study (≤100 pM), the majority of liposomes have a single PFO oligomer at the time of insertion and pore opening. It is important to stress that the formation of a second PFO dimer on the membrane does not necessarily lead to two independent pores as it could in principle join the end of the existing oligomer before membrane insertion. As such, this calculation should be taken as an upper limit for the probability of observing two independently growing structures on the liposome membrane.

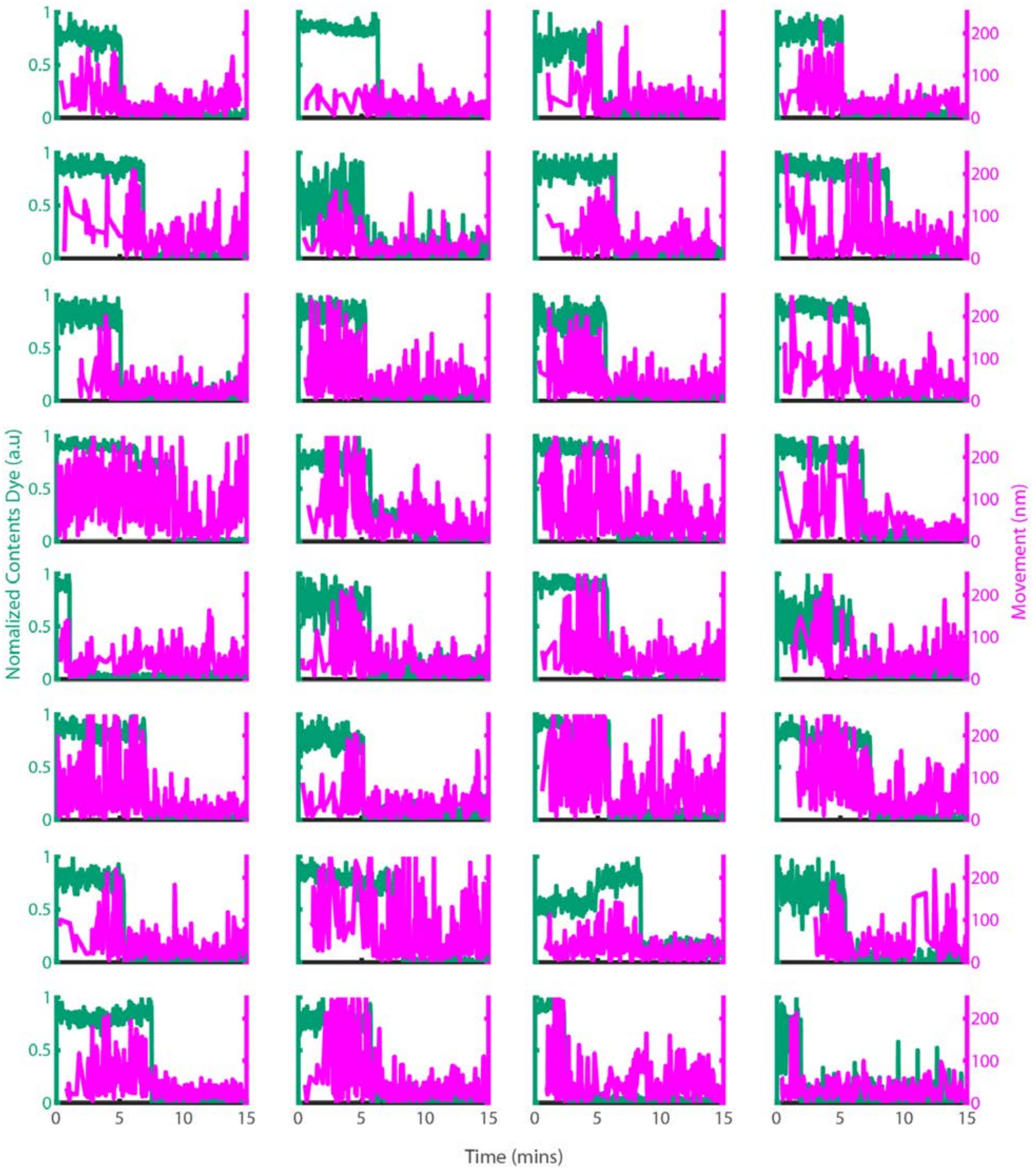


**Figure 7–Figure Supplement 1. Example traces of PFO oligomer movement on liposomes before and after poration.** Dye-loaded liposomes were imaged in the presence of 500 pM AF647-PFO by dual-colour TIRF microscopy (1 frame every 2 seconds, 50 ms exposure time, 2 mW 488 nm laser, 75 mW 640 nm laser). Pore opening was detected by release of the content dye while the x/y-positions of the PFO oligomer assembling on the liposome were tracked by point-spread function fitting. The movement of the AF647-PFO signal was calculated as the Cartesian distance between x/y-positions in subsequent frames. Shown are example traces recorded at single liposomes of the content dye signal intensity

(blue-green) and the corresponding movement of the position of the AF647-PFO signal between frames (magenta).

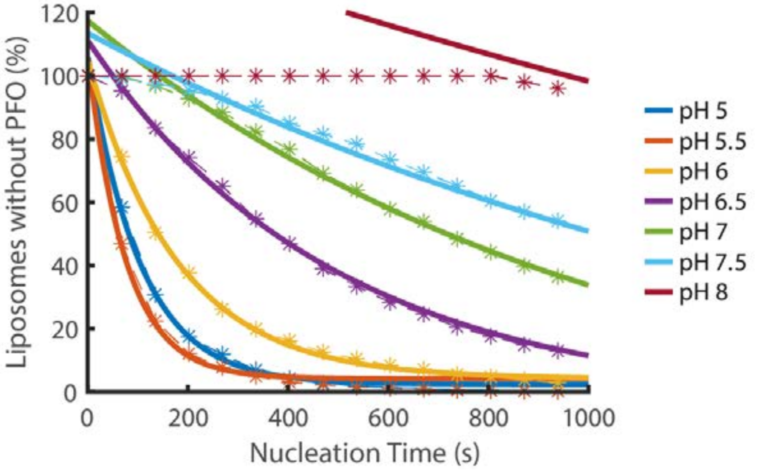


**Figure 8–Figure Supplement 1. Nucleation kinetics decrease with increasing pH.** Distributions of nucleation times measured at various pH in the presence of 200 pM AF647-PFO. Experimental data represented by dashed lines with stars. Each concentration is an average of at least three experiments. Exponential decay functions (solid lines) were fitted to the experimental data to determine the nucleation rate at each concentration.

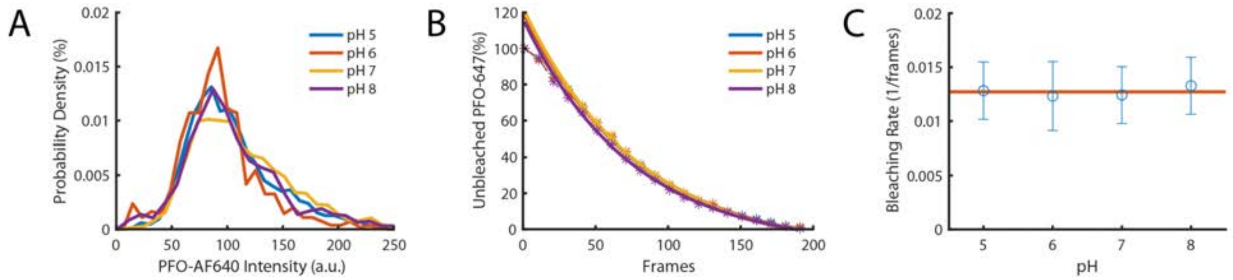


**Figure 8–Figure Supplement 2. The intensity of PFO-AF647 is independent of pH. (A)** The distribution of single-molecule intensities measured using single-molecule photobleaching for different pH buffers (pH 5, pH 6, pH 7 and pH 8). Data comprises 10 fields of view for each concentration. **(B)** The distributions of single molecule bleaching times obtained at different pH values are overlaid with exponential fits. **(C)** The corresponding bleaching rates from the exponent of the fits in panel B. The red line denotes the mean bleaching rate for all pH values.

### Appendix to “Single-molecule analysis of the entire perfringolysin O pore formation pathway”.

Conall Mc Guinness, James C. Walsh, Charles Bayly-Jones, Michelle A. Dunstone, Michelle Christie, Craig J. Morton, Michael W. Parker, Till Böcking

#### PFO amino acid sequence

The amino acid sequence of the His-tagged PFO C459A protein used in this study is shown below. The N-terminal hexahistidine tag followed by a thrombin cleavage site (underlined) are highlighted in blue. This construct for recombinant PFO expression in *E. coli* used in this study is missing the N-terminal signal peptide sequence (1-28) of the PFO precursor that is cleaved off during the biosynthesis of the functional protein in *Clostridium perfringens*. The numbering of the PFO residues is according to the full-length precursor protein (i.e. the construct for recombinant expression starts at K29).

```
MGSSHHHHHH SSGLVPRGSH MKDITDKNQS IDSGISSLSY NRNEVLASNG DKIESFVPKE   60
GKKAGNKFIV VERQKRSLTT SPVDISIIDS VNDRTYPGAL QLADKAFVEN RPTILMVKRK  120
PININIDLPG LKGENSIKVD DPTYGKVSGA IDELVSKWNE KYSSTHTLPA RTQYSESMVY  180
SKSQISSALN VNAKVLENSL GVDFNAVANN EKKVMILAYK QIFYTVSADL PKNPSDLFDD  240
SVTFNDLKQK GVSNEAPPLM VSNVAYGRTI YVKLETTSSS KDVQAAFKAL IKNTDIKNSQ  300
QYKDIYENSS FTAVVLGGDA QEHNKVVTKD FDEIRKVIKD NATFSTKNPA YPISYTSVFL  360
KDNSVAAVHN KTDYIETTST EYSKGKINLD HSGAYVAQFE VAWDEVSYDK EGNEVLTHKT  420
WDGNYQDKTA HYSTVIPLEA NARNIRIKAR EATGLAWEWW RDVISEYDVP LTNNINVSIW  480
GTTLYPGSSI TYN                                                     493
```

#### Mathematical analysis of PFO pore formation

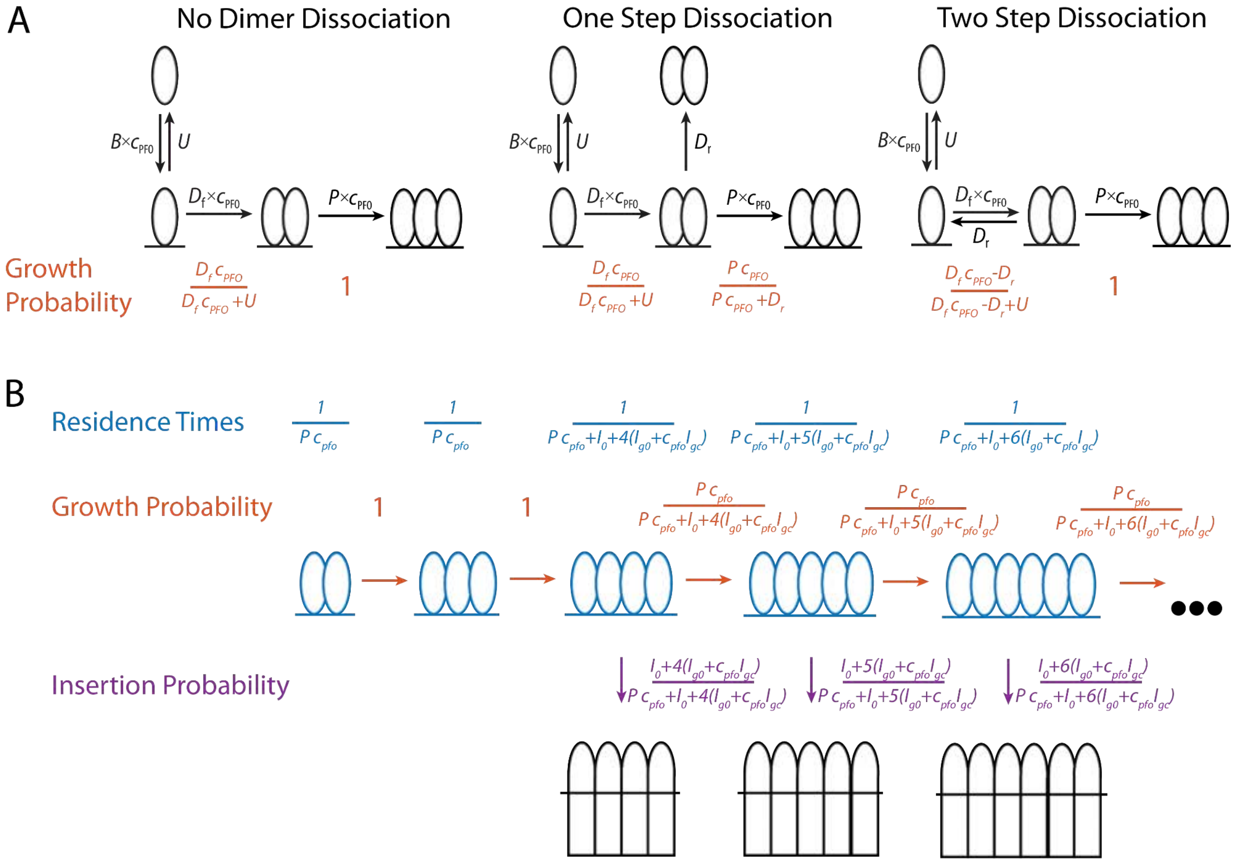


**Appendix Figure A1 Mathematical modelling of PFO. (A)** Schematics of the nucleation pathway for different scenarios of dimer dissociation **(B)** The residence times, and transition probabilities, of each state in the pore formation pathway. These are used to calculate the distribution of the number of subunits in the prepore at the time of insertion as well as the nucleation to insertion times.

**Nucleation analysis**

In Figure 3F, the dimer is shown to only be metastable, with a half-life of 13 minutes, although the pathway of dimer dissociation is not identified. The two potential methods of dimer dissociation, the dimer falling directly off the membrane in one step, or a two-step process, where the dimer falls apart into two membrane monomers which can then individually dissociate are shown in Figure A1 in the middle and right-hand schematics respectively. Here we calculate the nucleation rates for these two scenarios and compare them to the scenario the dimer was irreversibly bound (Figure A1A Left).

The probability that a monomer will become a dimer and a dimer will become a trimer is shown in orange below each transition in Figure A1A. In general, the probability that a single molecule will grow rather than fall off the membrane is given by the rate of growth divided by the sum of all rates out of that state. The one exception to this is in the two-step model where the rate of dimerisation is substituted by an effective rate of dimerisation ($D_f\, C_{pfo}$-$D_r$).

The nucleation rate can be modelled as the rate of monomer binding to the membrane ($B\, c_{pfo}$) multiplied by the probability that the monomer forms a dimer then forms a trimer. For no dimer dissociation this gives a rate of:

$$Nucleation\,rate\ =\ B\, c_{pfo}\frac{D_f\, c_{pfo}}{D_f\, c_{pfo}+U}$$

For a single step dimer dissociation the nucleation rate is

$$Nucleation\,rate\ =\ B\, c_{pfo}\frac{D_f\, c_{pfo}}{D_f\, c_{pfo}+U}\frac{P\, c_{pfo}}{P\, c_{pfo}+D_r}$$

For a two-step dimer dissociation the nucleation rate is

$$Nucleation\,rate\ =\ B\, c_{pfo}\frac{D_f\, c_{pfo}-D_r}{D_f\, c_{pfo}-D_r+U}$$

These equations are plotted in Figure 4–Figure Supplement 2B. The difference between the two release scenarios and the no dissociation scenario are shown in Figure 4–Figure Supplement 2C. For values above 100 pM the difference is less than 5%, suggesting that the dimer off rate is not a significant factor during nucleation.

**Insertion analysis**

Following nucleation PFO oligomerises into an arc (or potentially a full ring) before inserting into a ring. At each oligomer size (in molecules), the pore formation process can be thought of as a choice, either the oligomer adds an additional monomer and grows, or it inserts to form a pore. The likelihood of each of these choices depends on how quickly the oligomer is growing in number of subunits, and how fast it is able to insert. The higher the solution concentration of PFO, the faster it will oligomerise, and the more likely it is to grow rather than insert. Conversely, the more subunits in the prepore, the faster it inserts (Figure 5C and Figure 6A).

Mathematically, the probability of whether a prepore oligomer will grow, or insert, is given by the rate of oligomerisation ($P\, c_{pfo}$), or insertion ($I_0$+n ($I_{g0}$+$c_{pfo}\, I_{gc}$)) respectively, divided by the sum of these two rates. This is shown schematically in Figure A1B with growth probabilities coloured in Orange and insertion probabilities shown in Purple.

Overall, the probability that a pore will insert with a given number of subunits is then given by the product of the probabilities that the arc grew for all sizes less than the number of subunits at insertion before inserting at that number of subunits. For example, the probability of a pore inserting as a 5mer is:

$$Prob(5) = 1 \times 1 \times \frac{P\,c_{pfo}}{P\,c_{pfo}+I_0+4(I_{g0}+c_{pfo}I_{gc})} \times \frac{I_0+5(I_{g0}+c_{pfo}I_{gc})}{P\,c_{pfo}+I_0+5(I_{g0}+c_{pfo}I_{gc})}$$

This can be generalised for any number of subunits, n as:

$$Prob(n) = \frac{I_0+n(I_{g0}+c_{pfo}I_{gc})}{P\,c_{pfo}+I_0+n(I_{g0}+c_{pfo}I_{gc})} \prod_{j=m}^{n-1} \frac{P\,c_{pfo}}{P\,c_{pfo}+I_0+j(I_{g0}+c_{pfo}I_{gc})}$$

$$= I_0^{m-n-1}(I_0 + n(I_{g0} + c_{pfo}I_{gc}))(P\,c_{pfo})^{n-m}/pch((I_0 + m\,(I_{g0} + c_{pfo}I_{gc}) + P\,c_{pfo})/(I_{g0} + c_{pfo}I_{gc}), n + 1 - m)$$

Where *m* is the minimum number of subunits for insertion (assumed to be 4) and *pch* is the pochhammer function.
The size distribution predicted by this equation for various concentrations is shown in Figure 6–Figure Supplement 2B. These distributions are directly comparable to the experimental distributions shown in Figure 6–Figure Supplement 2A.

Any oligomers that reach 35 subunits are assumed to form complete rings and cannot oligomerise any further, only insert. The percentage of oligomers that form a complete ring is shown in Figure 6D. The mean of the number of subunits at insertion distribution is plotted as a function of concentration in Figure 6C where it is also compared to the mean of experimental data.

**Nucleation to insertion time**

During the oligomerisation process, we can calculate the average amount of time that an oligomer will spend with a given number of subunits before deciding to either grow or insert. The average total time from nucleation to insertion for a pore with a given number of subunits can then be calculated by summing these times.

The mean residence times for each state is given by the inverse of the sum of rates out of that state. This is shown schematically in blue in Figure A1B. The average time take for an oligomer to reach a certain number of subunits is then given by the sum of the times it spends in each size up to that number of subunits:

$$Growth\,Time(n) = \frac{m-2}{Pc_{pfo}} + \sum_{j=m}^{n} \frac{1}{Pc_{pfo}+I_0+j(I_{g0}+c_{pfo}I_{gc})}$$

To generate a mean nucleation to insertion time for all pores, the mean insertion time for each number of subunits is weighted by the probability of insertion occurring with that number of subunits (calculated in the Insertion Analysis section). A plot of the mean nucleation to insertion time for all pores as a function of concentration is shown in Figure 6E.

**The effect of pH on nucleation**

The pH dependence of the PFO oligomerisation rate shown in Figure 8C was heuristically fit with a straight line and taken as:

$P([pH]) = 0.243 - 0.03[pH]$

The pH dependence could result from either the membrane-binding kinetics of monomeric PFO, or the lateral interaction between membrane-bound PFO monomers being affected by changes in pH.

In the scenario where pH affects membrane binding (or release) then both the membrane binding and polymerisation interactions are affected. The value of single molecule binding (*B*) was scaled to change proportionately to the change in polymerisation:

$$B([pH]) = \frac{B([pH]=7)}{D_f([pH]=7)} D_f([pH]) = 0.95 - 0.12[pH]$$

Note that since single molecule binding to liposomes is transient ($U >> D\, c_{pfo}$ ) either single molecule binding (*B*) or unbinding (*U*) can be scaled and give the same results.

These two rates were then substituted into the nucleation rate equation from the section Nucleation Analysis:

$$Nucleation\, rate([pH]) = B([pH])\, c_{pfo} \frac{P([pH])\, c_{pfo}}{P([pH])\, c_{pfo} + U}$$

This line is plotted in purple in Figure 8D.

In the case that pH only affects PFO-PFO lateral interactions, then only the polymerisation rate would be affected so the equation would then be:

$$Nucleation\, rate([pH]) = B\, c_{pfo} \frac{P([pH])\, c_{pfo}}{P([pH])\, c_{pfo} + U}$$

This line is plotted in yellow in Figure 8D.

**The Effect of pH on growth and insertion**

The dependence of insertion kinetics was modelled by substituting the heuristic oligomerisation rate dependence fit from Figure 8C (See “The Effect of pH on Nucleation”):

$$P([pH]) = 0.243 - 0.03[pH]$$

Into the equations for insertion distribution (see “Insertion Analysis”) and nucleation to insertion time (see “Nucleation to Insertion Time”). The concentrations were then set to equal experiments (200 pM) and rates were plot as a function of pH to generate curves shown in Figure 8F-H

#### Error calculation of parameter estimates

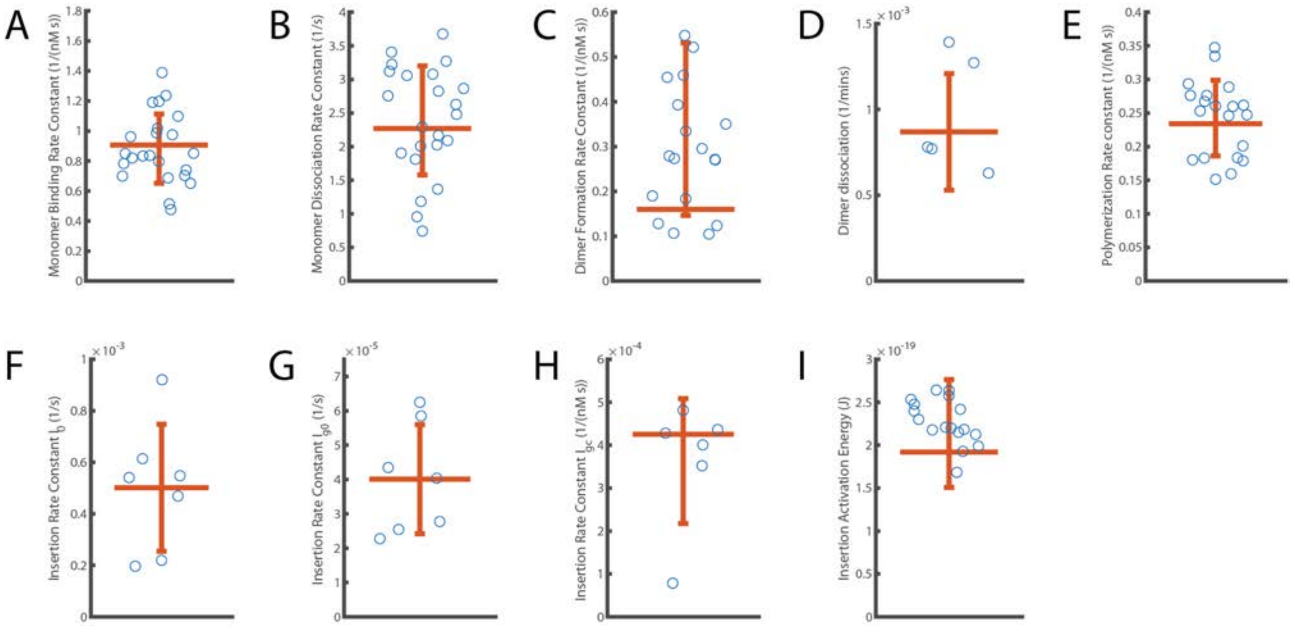


**Appendix Figure A2.** Distribution of independent measurements for each parameter. The error bar displays the standard deviation of measurements around the mean. The horizontal orange line shows the aggregated fit value reported for each parameter. **(A)** Monomer binding rate constant (*B*; from Figure 2). **(B)** Monomer unbinding rate (*U*; from to Figure 2). **(C)** Dimer formation rate constant ($D_f$; from Figure 3). **(D)** Dimer dissociation rate ($D_r$; from Figure 3). **(E)** Oligomerisation rate constant (*P*; from Figure 4). **(F-H)** Components describing the overall kinetics of insertion during continuous oligomer growth: insertion rate ($I_0$, shown in panel F), oligomer subunit number-dependent insertion rate ($I_{g0}$, shown in panel G) and PFO concentration-dependent insertion rate ($I_{gc}$, shown in panel H) (from Figures 5 and 6). **(I)** Activation energy for insertion, i.e. transition from the prepore state to the open pore state ($E_a$; from Figure 5).

#### Single-Molecule Photobleaching for Intensity Calibration

A single-molecule photobleaching experiment was used to calculate conversion factors to convert measured fluorescent intensities to numbers of bound molecules. Measuring the photobleaching rate also governed how many frames were acquired before bleaching becomes significant.

The protocol for photobleaching was as follows:

1) 25mm round Coverslips were cleaned by sonication in ethanol, water, 1M NaOH then water again for 15 minutes each before being blow-dried with nitrogen.
2) A coverslip was exposed to an air plasma using a plasma cleaner (Harrick Plasma) before being placed in a Chamlide chamber to prevent liquid from running off the edge of the slide.
3) 1 mL of the fluorescently labelled PFO at 50 pM was added to the coverslip and left to bind for 5 minutes.
4) The supernatant was removed and the sample was washed with 1 mL clean buffer to remove unbound molecules.
5) Wash buffer was then discarded and replaced with fresh wash buffer.
6) The sample was then imaged on the microscope (Appendix Figure A3). A photobleaching image stack was collected by exposing a field of view with the same laser power setting used during the actual experiment but at four times the exposure time (200 ms).
7) 200 frames were imaged so that approximately 90% of fluorophores were bleached.
8) Five fields of view were imaged to measure variability within the sample and consistency of analysis.

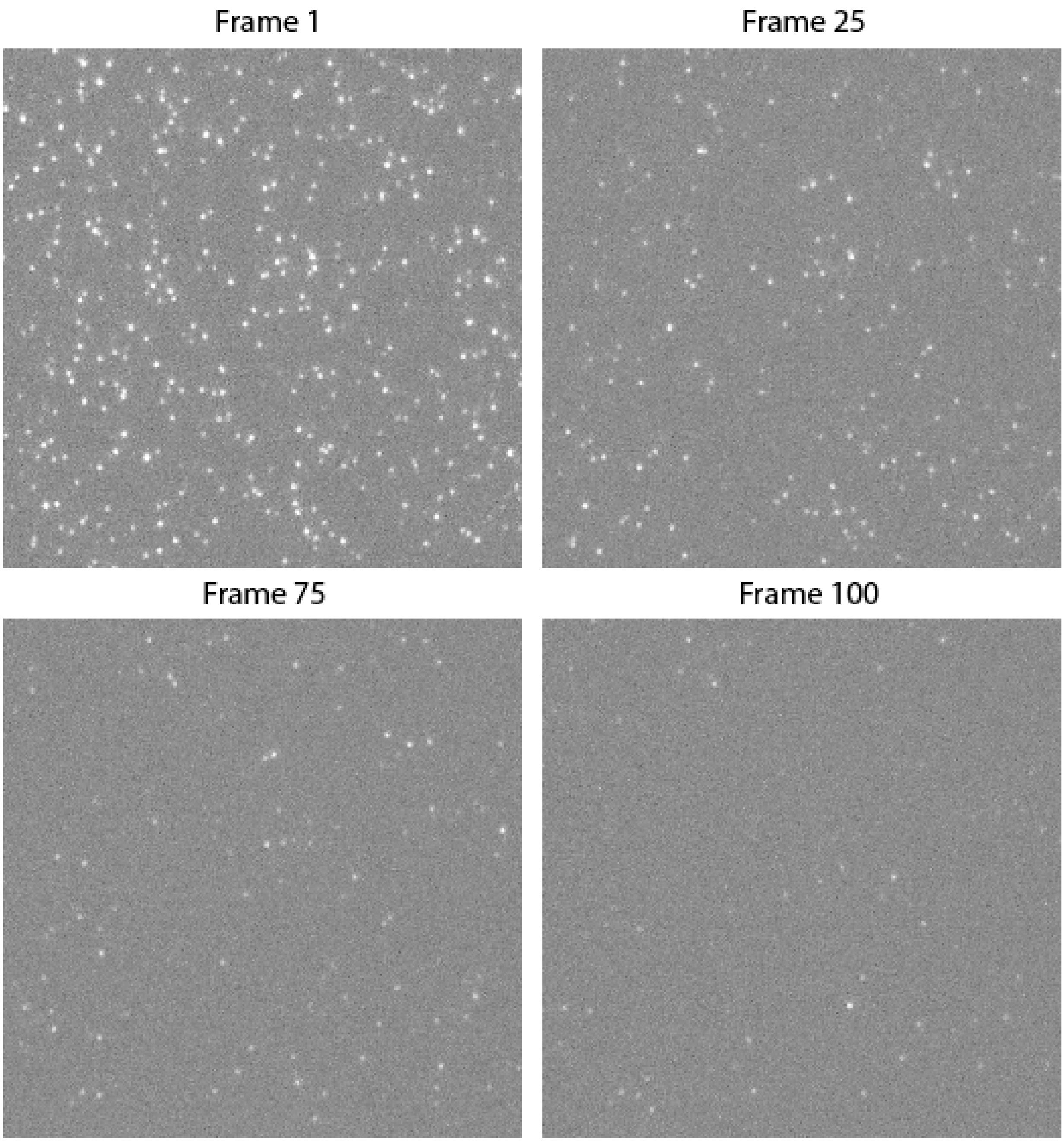


**Appendix Figure A3. Montage of Single-Molecule Photobleaching Image Stack.** Single PFO particles, immobilised on a coverslip, bleach as more frames are imaged. The montage shows the 512x512 pixel centre of a single field of view. The complete image is 2048x2048 and in total 5 fields of view were imaged.

##### Photobleaching Image Analysis

Image stacks are analysed using the JIM software package developed in our lab. This software is open source, continuously maintained, and freely available at https://github.com/lilbutsa/JIM-Immobilized-Microscopy-Suite. The documentation for JIM contains multiple tutorials including a step by step guide for collecting and analysing single-molecule photobleaching. The documentation is available at https://docs.google.com/document/d/12frP6jp74eiycXxY8knR_qb27ui7Ia-RNMBuAuAHRFs/edit

Traces are generated for photobleaching image stacks by detecting regions of interest (ROIs) from the first 10 frames of the experiment. These traces are then analysed using the step fitting program (part of the JIM package) which uses change point analysis to heuristically determine whether or not a step occurs in a trace. An in-depth description of the step fitting program is available in the relevant section of the documentation.

After step fitting, three filters are applied to the step traces to determine which traces to include in the analysis. These filters are as follows :
- first step probability: 0.5
- ratio between the second and first intensity level: 0.25 (at least 75% of the signal is lost)
- probability of more steps: 0.999 (exclude traces with additional steps)

The photobleaching rate is obtained by fitting an exponential decay function to the distribution of bleaching times (Figure 2–Figure Supplement 3A/B). The single-molecule intensity is obtained from the mean of a log-normal curve fitted to the distribution of step heights (Figure 2D).
